# Charting Improvements in US Registry HLA Typing Ambiguity Using a Typing Resolution Score

**DOI:** 10.1101/050443

**Authors:** Vanja Paunić, Loren Gragert, Joel Schneider, Carlheinz Müller, Martin Maiers

## Abstract

Unrelated stem cell registries have been collecting HLA typing of volunteer bone marrow donors for over 25 years. Donor selection for hematopoietic stem cell transplantation is based primarily on matching the alleles of donors and patients at five polymorphic HLA loci. As HLA typing technologies have continually advanced since the beginnings of stem cell transplantation, registries have accrued typings of varied HLA typing ambiguity. We present a new *typing resolution score*, based on the likelihood of self-match, that allows the systematic comparison of HLA typings across different methods, data sets and populations. We apply the typing resolution score to chart improvement in HLA typing within the Be The Match Registry of the United States from the initiation of DNA-based HLA typing to the current state of high-resolution typing using next-generation sequencing technologies. In addition, we present a publicly available online tool for evaluation of any given HLA typing. This *typing resolution score* objectively evaluates HLA typing methods and can help define standards for acceptable recruitment HLA typing.

## 1.1 Introduction

The Human Leukocyte Antigen (HLA) genes are the most polymorphic genes in the human genome, with more than 12,000 variations (alleles) identified to date. As an immune compatibility gene system, HLA plays a critical role in hematopoietic stem cell transplant (HSCT), where the degree of HLA matching is the major determinant of outcome [1,2]. Most patients in need of HSCT identify their donor through unrelated stem-cell registries worldwide due to the unavailability of matched related stem-cell donors in 70% of cases. Initially, HLA specificities were assessed by serology and donors were selected based on matching for these antigens. In the last decade, outcomes studies have revealed that allele-level matching leads to superior outcome relative to serologic antigen matching [3]. Unfortunately, identification of allele-matched donors in registries is hindered by the accrual of large numbers of donors with legacy HLA typings where the exact alleles present were not determined.

Exact allele-level HLA typing used to be costly and laborious with the large and rapidly growing number of described HLA alleles. The characterization of HLA alleles in patients seeking HSCT is generally performed by allele-level HLA typing at four or more HLA loci. If familial typing is available, HLA haplotype phasing may also be assigned. In contrast, HLA typing of donors in unrelated donor registries is often considerably more ambiguous, consisting of lists of possible HLA alleles that the donor might have. Ambiguous allele assignments are produced due to either failure to interrogate all polymorphic positions or lack of phase between polymorphisms within a locus because of diploid sequence reads (or both). DNA-based methods identify HLA alleles by interrogating the nuclear DNA sequence and can result in different levels of ambiguity depending on typing methodology or test kits used. Some of the widely used DNA-based HLA typing methods are sequence-specific oligonucleotide (SSO) hybridization, sequence-specific primer (SSP) amplification and Sanger sequence-based typing (SBT).

To overcome the challenge of HLA typing ambiguity in identifying allele-matched donors, unrelated stem-cell registries often apply statistical methods to predict which donors are most likely to be allele-level matched [4–7]. The high-resolution HLA haplotype frequencies that enable these predictions are calculated on large samples from diverse ethnic groups. High linkage disequilibrium allows for HLA typing information at one locus to inform predictions at other HLA loci. One major factor impacting the performance of HLA imputation is the ambiguity content of the input HLA typing.

We recently demonstrated that Shannon’s entropy, a measure originally used in information theory, successfully quantifies the ambiguity content in HLA typing and can be used to objectively compare different typing methods to each other [8]. Our results showed that single-pass sequence-based typing (SBT) reported in genotype list format gives high certainty in allele prediction across all populations. Our results were supported in [9] where the authors advocate the use of a one-step DNA sequencing strategy to HLA type hematopoietic stem cell donors at recruitment.

In this paper we expand on our previous efforts to quantify HLA typing ambiguity [8]. We first describe a more intuitive measure for quantifying typing ambiguity that is comparable across different system. We refer to it as Typing resolution score (TRS). We then use TRS in order to analyze the HLA typing data in the Be The Match Registry charting the improvements in the Registry across populations and across the history of registry from the inception of DNA-based typing in 1993. We aim to measure the ambiguity in HLA typing in the Registry at the locus level, and at the levels of unphased and phased HLA genotypes. The locus-wise analysis of ambiguity primarily reflects aspects of typing technologies and methods. The unphased genotype-level analysis reflects population aspects of donor and patient matching relevant for search coordinators, since donor and patient matching is currently performed at the level of unphased HLA genotypes. There is evidence, however, that phase matching for fully allele-matched unrelated donors and patients may improve outcomes post-transplant [10]. Therefore, we also aim to evaluate the amount of ambiguity in phased HLA genotype assignment in the Registry.

## 1.2 Materials and Methods

### 1.2.1 Dataset

Be The Match Registry, operated by the National Marrow Donor Program (NMDP), began HLA typing of unrelated donors in 1987. We used the HLA genotypes of 15,697,842 adult donors and 436,437 cord blood units. We restricted the analysis to the samples that are at the minimum typed at HLA-A, -B, and -DRB1. To compute all possible genotypes from an ambiguous HLA typing [4] as a part of computing the typing resolution score, we used high-resolution haplotype frequencies generated from unrelated donors from the NMDP database for five population categories described in [11]. These categories are created from the self-described race and ethnicity backgrounds recorded at donor registration. It should be stressed that the typing resolution score depends on the population, as population-specific HLA haplotype frequencies are used in the computation of the score. The summary of the Registry data used in this study is shown in the Table 1.

**Table 1.** **Study Data**. Be The Match Registry/NMDP HLA genotypes used in the study, by the race/ethnicity group and the sample type.

| Broad population | Cord blood units (CBU) | Donors |
| --- | --- | --- |
| African American (AFA) | 26,839 | 965,087 |
| Asian/Pacific Islander (API) | 27,653 | 990,356 |
| European American (CAU) | 324,632 | 12,305,015 |
| Hispanic (HIS) | 56,379 | 1,307,072 |
| Native American (NAM) | 934 | 130,312 |
| **Total** | **436,437** | **15,697,842** |

The analysis is performed for five HLA loci, HLA-A, -C, -B, -DRB1, -DQB1, at the antigen recognition site (ARS) exons only level of allele resolution.

### 1.2.2 Typing resolution score

We define typing resolution score (TRS) as:

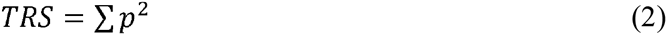

where *p* is a set of normalized frequencies of distinguishable types, in our case a set of allele or genotype frequencies. The typing resolution score has several attractive mathematical properties. It is bounded on [0, 1] interval, where 0 indicates maximum ambiguity, and 1 indicates no ambiguity in an HLA typing. As such, TRS can be used to compare typings generated across different systems, as its maximum value no longer depends on the length of the imputation output (as was the case with the entropy-based measure). This becomes even more relevant as we hope that the score proposed here can help define acceptable typing at the donor recruitment.

### 1.2.3 Phased genotype typing resolution score

Given an ambiguous typing, HLA genotype imputation methods [4] resolve this ambiguity by generating a set of phased genotypes that could be expanded from the typing. To obtain a frequency for each phased genotype, the methods use HLA population haplotype frequencies [11] and calculate the frequency using equation *δ* * *f_1_*\* *f*_2_, where *f*_1_ and *f*_2_ are frequencies of two haplotypes that make up the phased genotype and *6* a constant that equals 1 in case of homozygous haplotypes, and equals 2 in case of heterozygous haplotypes. A set of all frequencies *p* is obtained by repeating the above calculation for each phased genotype that could be obtained from the HLA typing. To compute a typing resolution score for phased genotypes, we use the obtained set of frequencies *p* in the Equation 2. An example of ambiguous typing, resolved phased genotypes and the associated frequencies, is shown in Table 2.

**Table 2.**
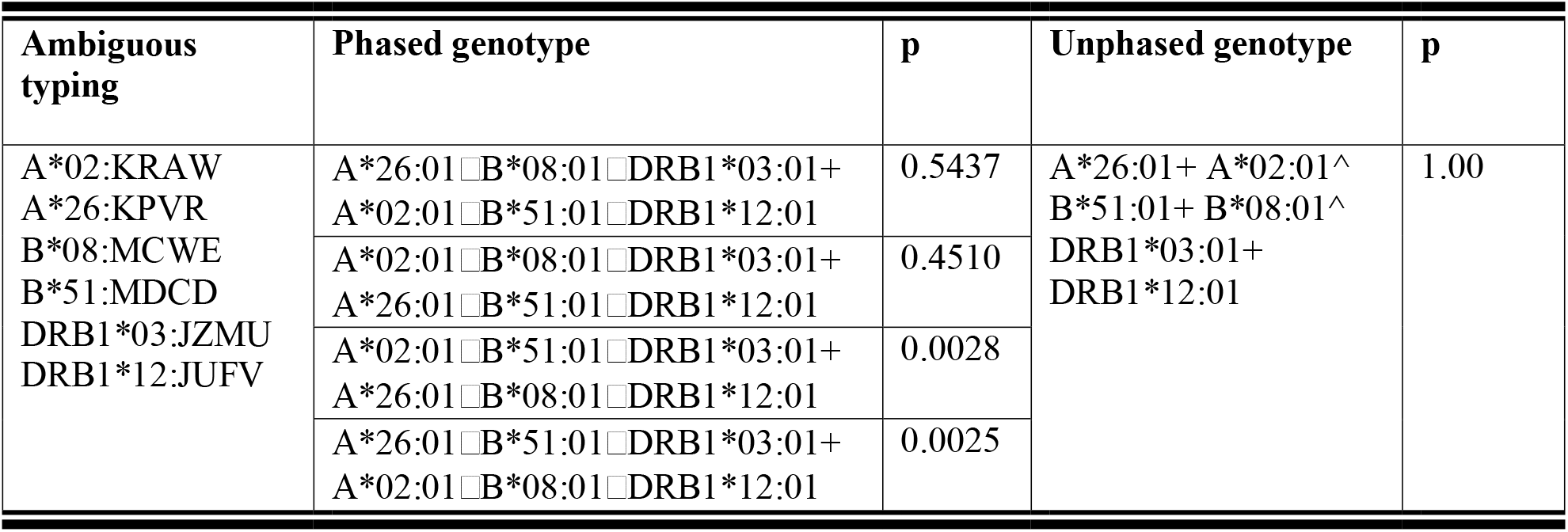
**An example of ambiguous HLA typing**. This table shows an example of ambiguous HLA typing as produced by a typing laboratory. The second and third columns show the result of HLA imputation applied to this HLA typing along with the assigned normalized frequencies p for each phased genotype. The fourth column shows all unphased genotypes that can be obtained from the imputed phased genotypes. In this example, only one unphased genotype can be obtained from the imputed phased genotypes, and its associated normalized frequency is equal to 1 (summation of all associated phased genotype frequencies). Note that the phased and unphased genotypes are represented in GL string format, a preferred data reporting format ambiguous data without loss of typing information [21].

### 1.2.4 Unphased genotype typing resolution score

In the context of matching between donor and patient HLA alleles for bone marrow transplant, we are only interested in resolving allele ambiguity, and not the phase ambiguity. The patient and the donor are considered a match for bone marrow transplant if their two alleles at a required HLA loci are the same, without regard to their chromosomal assignment, that is, at the level of unphased genotype. The frequency of unphased genotype *g* can be obtained from the phased genotype frequencies, by summing over the frequencies of all phased genotypes that could be obtained from the genotype *g*. This can be formally written as:

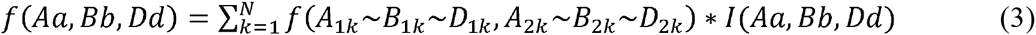

where *N* is the number of phased genotypes generated from the HLA typing and *I*(*Aa, Bb, Dd*) is an indicator function that takes value of 1 when the phased genotype *A*_1*k*_~*B*_1*k*_~*D*_1*k*_, *A*_2*k*_~*B*_2*k*_~*D*_2*k*_ can be collapsed into unphased genotype *Aa, Bb, Dd* and the value of 0 otherwise. The equation is written for three-loci genotypes only for the brevity of notation, but it can be applied to any number of loci. In this paper, we apply it to genotypes consisting of five loci, HLA-A, -C, -B, -DRB1, and -DQB1.

HLA ambiguity resolution to the level of unphased genotypes, rather than phased genotypes, removes a certain level of ambiguity from the typing, since often a given unphased genotype can be made up of more than one combination of known phased genotypes. A set of all frequencies *p* is obtained by computing the frequency for each unique unphased genotype that could be obtained from a given HLA typing. To compute typing resolution score for unphased genotypes, we use the obtained set of frequencies *p* in the Equation 2. An example of ambiguous typing, resolved unphased genotypes and the associated frequencies, is shown in Table 2.

### 1.2.5 Locus typing resolution score

To compute locus-wise typing resolution score we again use the output of HLA imputation for the given ambiguous HLA typing, that is, all possible phased genotypes and their frequencies. We then calculate frequencies of each unique allele pair on a given locus by summing over all frequencies of phased genotypes generated for the given typing, that contain the given allele pair, that is

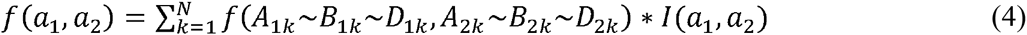

where *N* is the number of generated phased genotypes and *I*(*a*_1_, *a*_2_) is an indicator function that takes value of 1 when the phased genotype *A*_1*k*_~*B*_1*k*_~*D*_1*k*_*, A*_2*k*_~*B*_2*k*_~*D*_2*k*_ contains allele pair *a*_1_*, a*_2_ and a value of 0 otherwise. A set of all frequencies *p* is obtained for all unique allele pairs that could be obtained from a given HLA typing, and used in Equation 2 to compute locus-wise typing resolution score for the given locus.

## 1.3 Results

### 1.3.1 Analysis of the current Registry

In this section we describe the results on the Registry data, which contains all HLA genotypes present in the Registry as of May 16, 2015.

Figure 1(a) shows the distribution of TRS scores across different population groups for the unphased HLA genotypes. The differences in ambiguity between populations in the Registry confirm the findings in our previous work obtained on the simulated data [8]. It shows that the median TRS for CAU population is above 0.9, drastically higher than for other populations. This is partly due to the largest representation of CAU samples in the Registry and better representation of the HLA types which leads to better HLA haplotype frequency estimates and therefore better reduction in uncertainty through imputation, but also, due to the efforts to improve typing methods to separate the alleles for the majority group (CAU).

**Figure 1.**
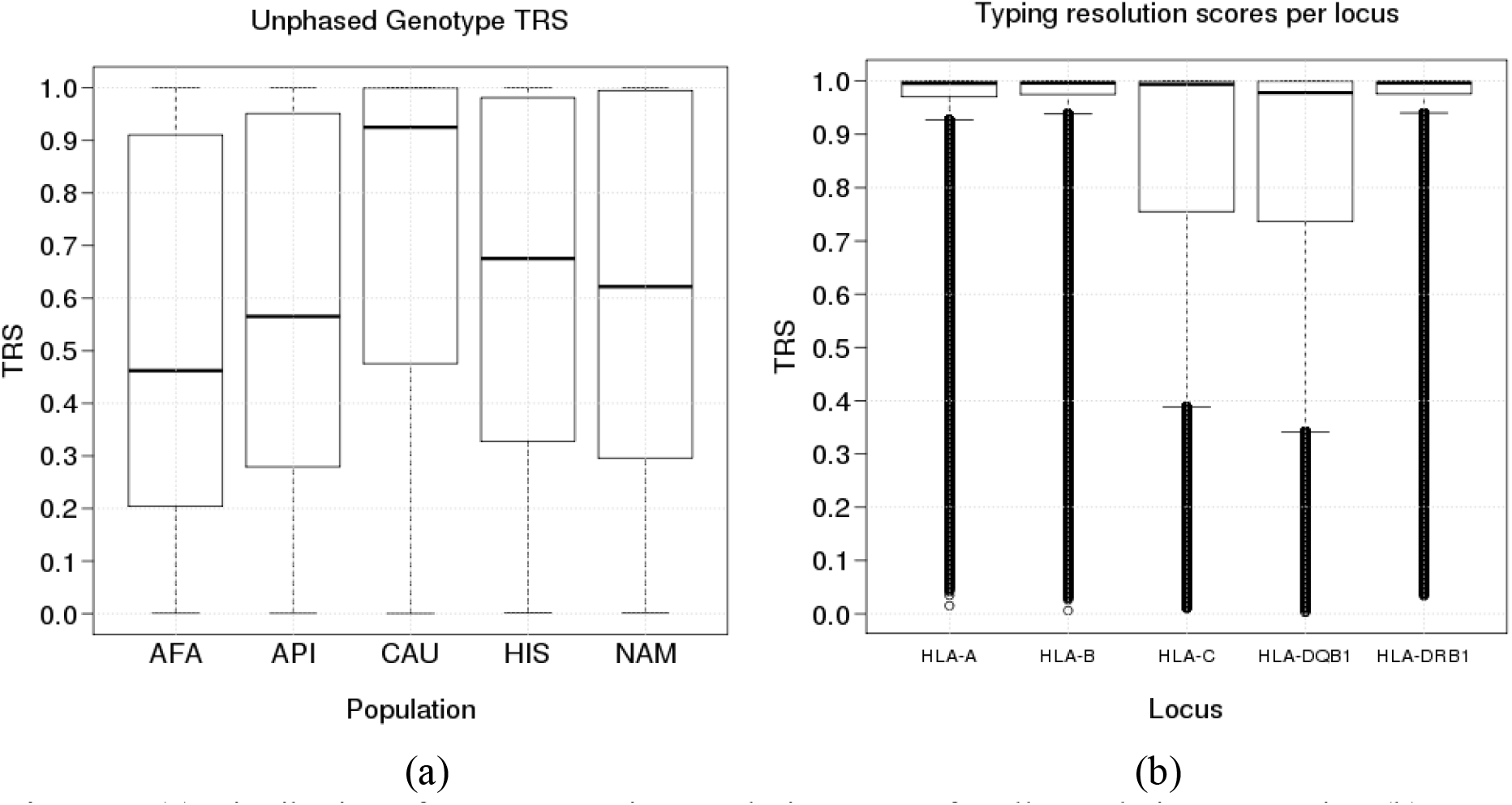
(a) Distribution of genotype typing resolution scores for all population categories. (b) Distribution of typing resolution scores for all five HLA loci.

Per-locus analysis of the current Registry is shown in the Figure 1(b) which shows the distribution of scores obtained for each HLA locus separately, and Figure 2 which shows the distribution of per-locus scores for each population separately. As expected, the loci that have been typed and accrued in the Registry longer, HLA-A, B and DRB1, have higher scores. Recruitment typing at these three loci fully replaced the previous requirement for typing at HLA-A and B only in 2001. On the other hand, the inclusion of HLA-C and HLA-DQB1 at recruitment at NMDP started only recently, in 2008 and 2010, respectively. However, due to confirmatory typing and re-typing efforts of historical data in the registry, some donors typed before this time have HLA-C and HLA-DQB1 loci typed. The differences of locus scores between populations are similar to our previous result on the overall genotype scores. Note that the median locus scores for all loci are higher than median scores for unphased genotypes. This is expected since evaluating ambiguity at each locus does not involve measuring ambiguity at other loci. The higher values at the locus level are also an artifact of the majority of the Registry being CAU samples, and their relatively low ambiguity, as shown in Figure 1(a).

**Figure 2.**
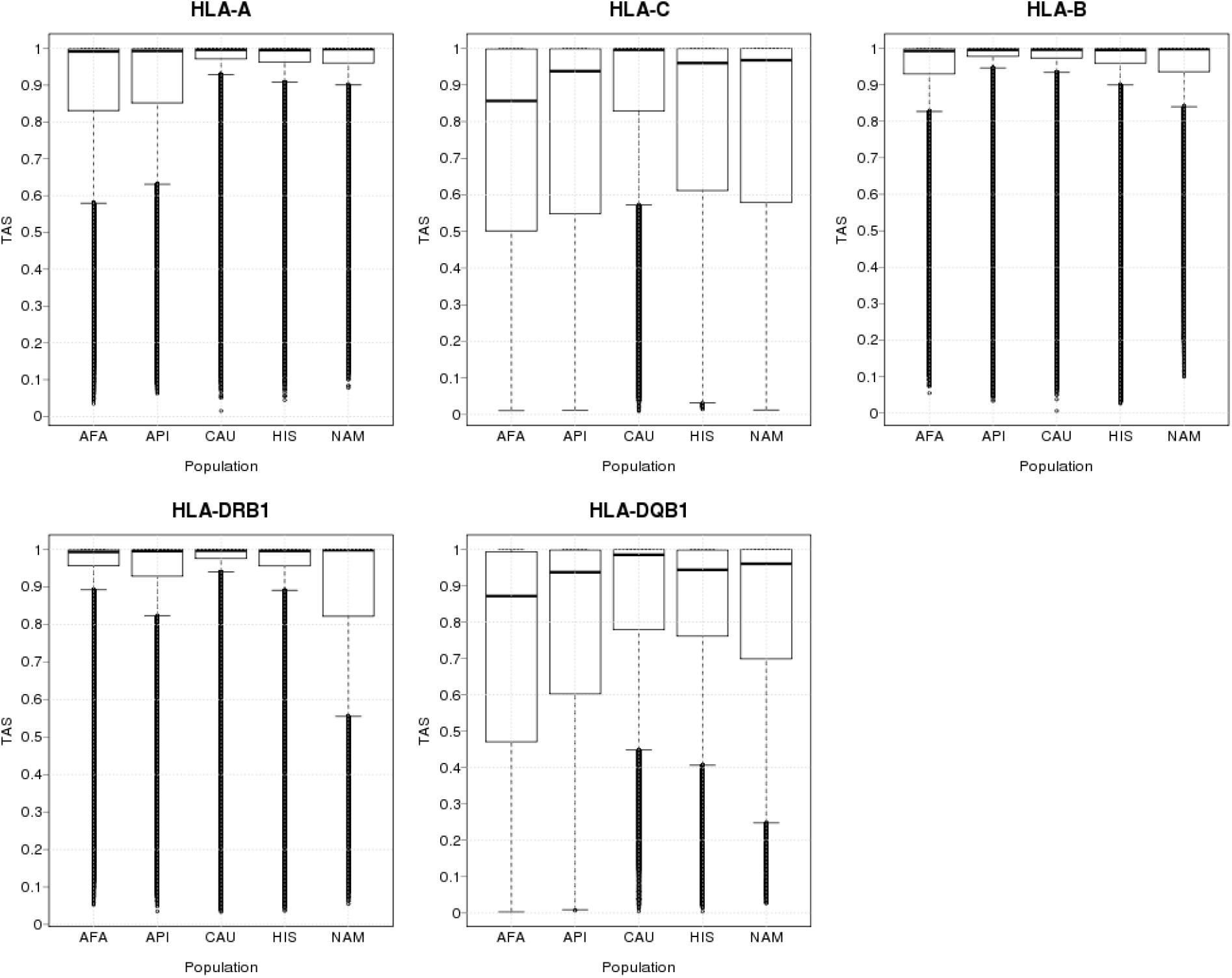
Distribution of typing resolution scores for each HLA locus and for each population category separately.

### 1.3.2 Analysis of the Registry over time

In this section we evaluate the ambiguity in the registry over time, starting in year 1993 when DNA-based typing began at NMDP. The number of donors in the Registry used in this study in each year is shown in Table 3.

**Table 3.** **Registry size over time**. The number of donors and cord blood units analyzed in this study for each year.

| Year | Number of donors |
| --- | --- |

|  | **and CBUs** |
| --- | --- |
| 1993 | 1,185,723 |
| 1994 | 1,594,452 |
| 1995 | 2,067,996 |
| 1996 | 2,695,432 |
| 1997 | 3,165,819 |
| 1998 | 3,655,773 |
| 1999 | 4,059,904 |
| 2000 | 4,499,744 |
| 2001 | 4,944,105 |
| 2002 | 5,364,097 |
| 2003 | 5,836,368 |
| 2004 | 6,349,905 |
| 2005 | 6,820,586 |
| 2006 | 7,348,392 |
| 2007 | 8,054,469 |
| 2008 | 8,765,366 |
| 2009 | 9,772,696 |
| 2010 | 10,865,890 |
| 2011 | 11,917,147 |
| 2012 | 13,014,923 |
| 2013 | 14,164,810 |
| 2014 | 15,588,773 |
| 2015 | 16,134,279 |

Figure 3 shows how the average typing resolution score changes over time at the unphased genotype level, as well as for each locus. Cumulative TRS for each year is obtained by averaging scores for all donors that are added to the registry that year or prior to that year, while recruitment TRS is obtained by averaging scores for only those donors that are added to the Registry in that year. While both recruitment and cumulative score follow an increasing trend over time, there is more variation in this trend in the early years of recruitment, partly due to smaller samples sizes in those years and more variability.

**Figure 3.**
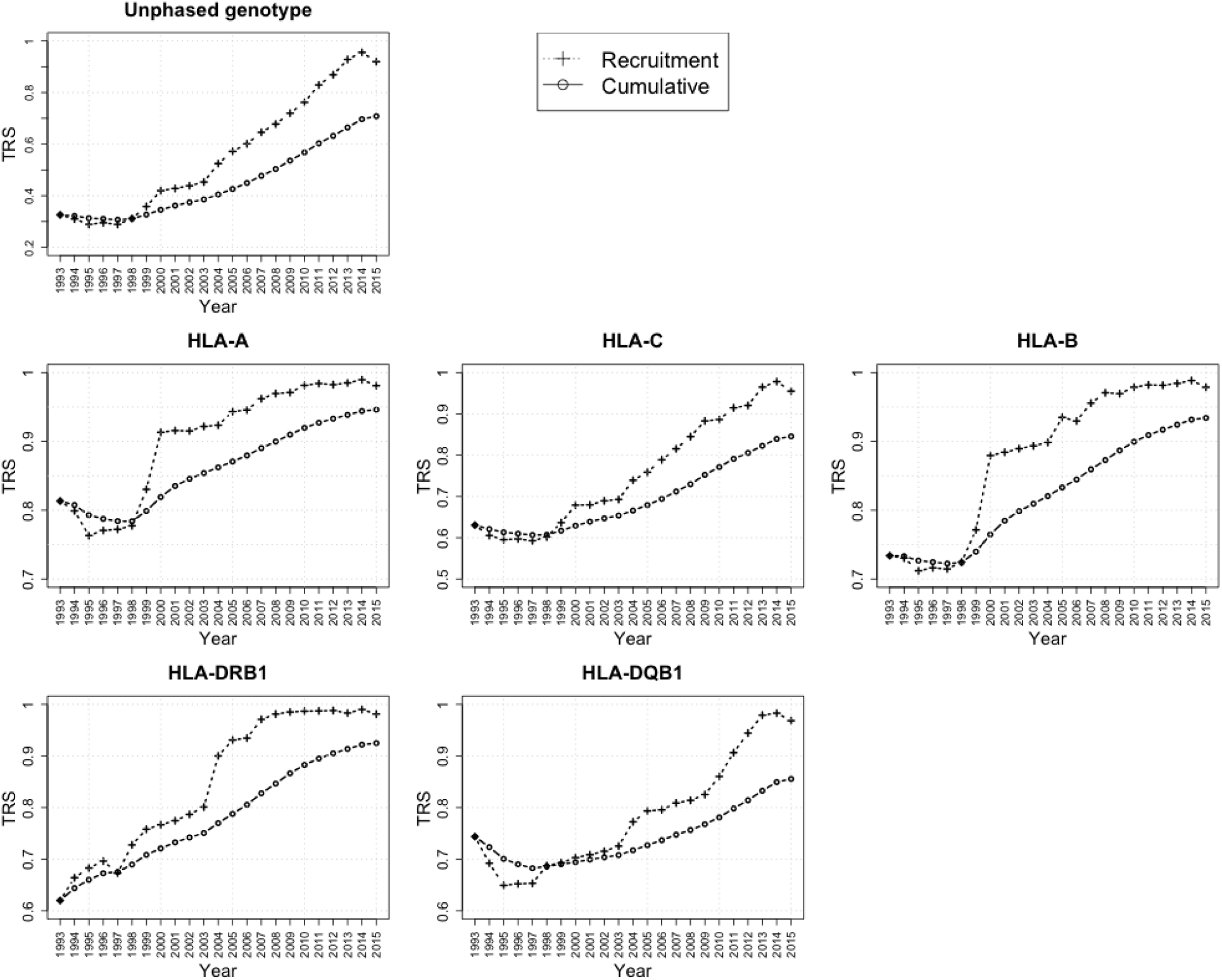
**Cumulative vs. recruitment score over time**. This figure shows cumulative vs. recruitment scores over time for overall genotypes and each HLA locus averaged over all samples. Cumulative score for each year is obtained by averaging scores for all samples that are added to the Registry that year or prior to that year, while recruitment score is obtained by averaging scores for only those samples that are added to the Registry in that year.

Figures 2s and 3s shows the trend of cumulative and recruitment TRS, respectively, over time for each population separately. Similar observations for each population can be seen here, with the CAU samples generally having the most rapid increase in scores, while African American donors have the lowest scores and a slower increase in scores over time. This trend demonstrates that oligo-based kits were tuned to reduce typing ambiguity primarily for majority race/ethnic groups, such as European American donors. Note that the trend of the score for HLA-DQB1 differs from that of the other loci in the earlier years. Since HLA-DQB1 was the last locus to be included in the requirement for typing at recruitment, so the earlier records included in this study may be skewed towards the higher resolution types coming from confirmatory typing and re-typing efforts of historical data in the Registry.

### 1.3.3 Phased genotypes TRS results

Here we describe the results of the evaluation of the ambiguity in the Registry at the phased genotype level. These results show, not only how much ambiguity the typings contain in the allele assignment, but also in the phase (chromosomal) assignment. HLA phase is currently not used in matching donors and patients in the Registry. There is evidence, however, that phase matching for fully allele-matched unrelated donors and patients may improve outcomes post-transplant. HLA haplotype mismatching was associated with a statistically significantly increased risk of severe acute GVHD and with lower risk of disease recurrence after the transplant [10]. Therefore, evaluating the ambiguity in phased HLA genotype assignment in the Registry is important.

Figure 4(a) shows the distribution of scores for the phased genotypes for all five broad populations. While the median trend across populations is similar to that of the unphased genotype in Figure 1(a), we see a significant decrease in the scores for all populations. The largest decrease in the median score when phase ambiguity is accounted for is seen in the European American population, where the median score drops from 0.91 to 0.58. The distribution of phased genotype scores vs. unphased genotype scores across all samples is shown in Figure 4(b). Average scores for overall genotype and phased genotypes for each population are shown in Table 1s. The difference between these two distributions reflects the amount of ambiguity remaining in the haplotype assignment of the HLA alleles, after the allelic ambiguity has been resolved. This is a demonstration that while the newer sequencing technologies have achieved a great reduction at allelic ambiguity at HLA loci, there still remains room for improvement when it comes to reducing the ambiguity in haplotype assignment for the HLA typings in the Registry.

**Figure 4.**
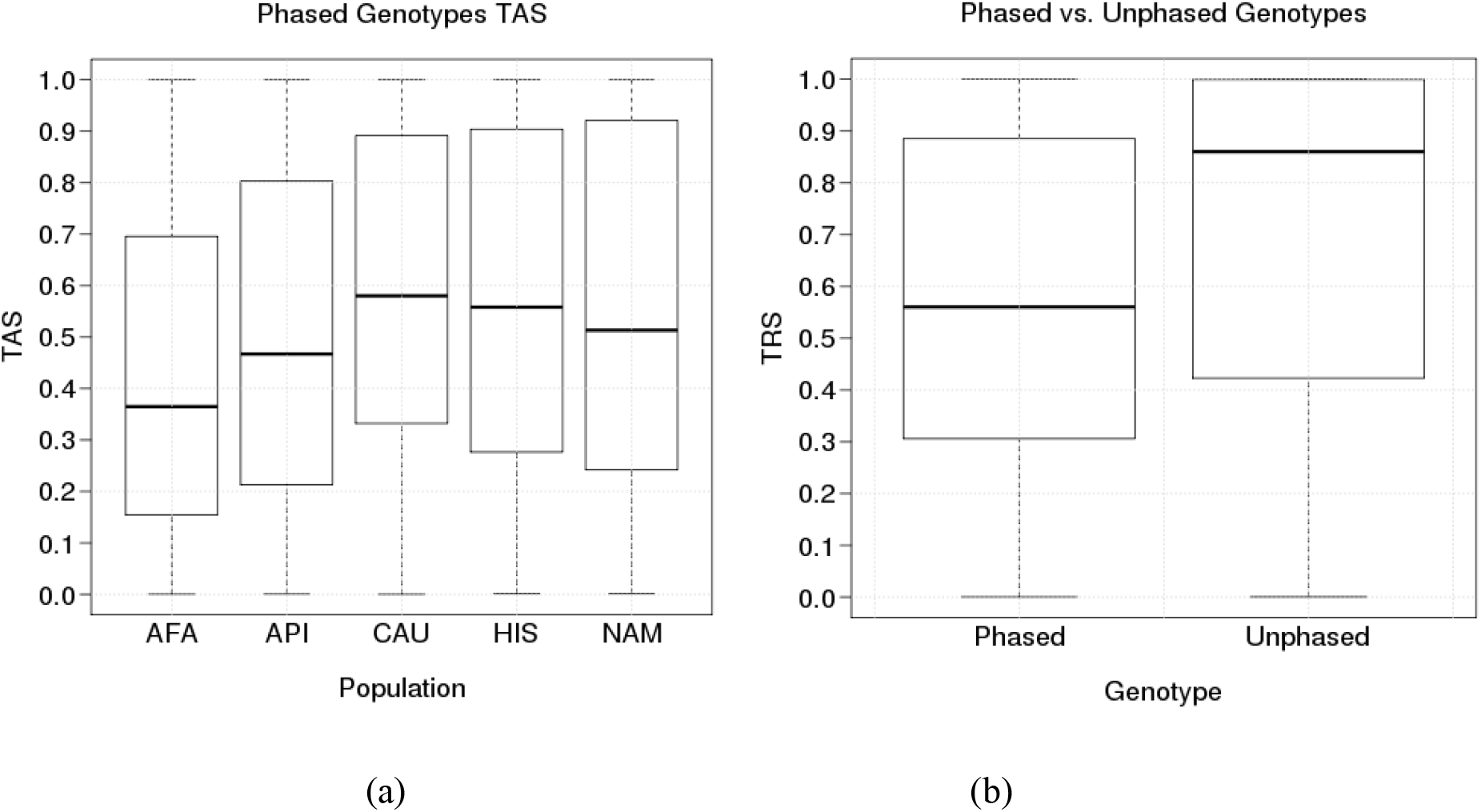
(a) Distribution of phased genotype typing resolution scores for all population categories. (b) Distribution of typing resolution score for phased genotypes vs. overall genotypes overall all samples.

Finally, Figure 5 shows the change of the phased ambiguity score over time for cumulative score, recruitment score, and the comparison between them, respectively. The overall trends in this Figure are similar to those for the unphased genotype score, while the values of the scores are lower.

**Figure 5.**
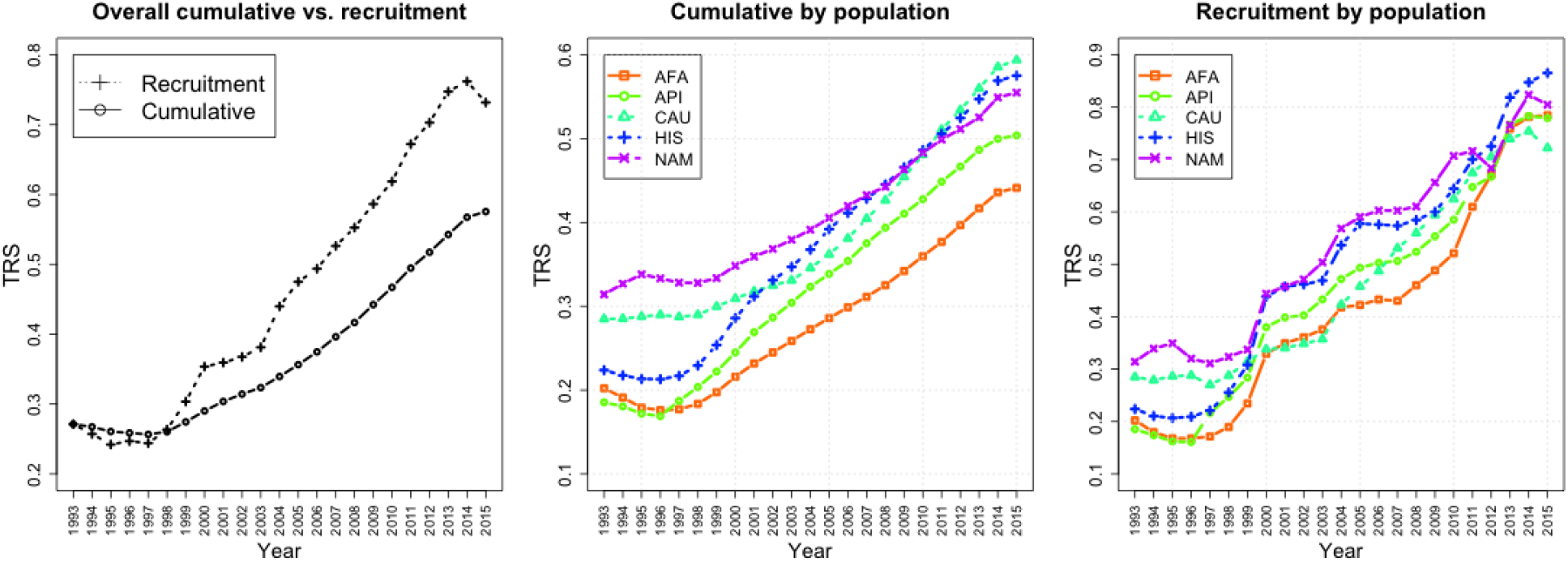
**Phased genotype typing resolution score over time**. This figure shows the change of phased cumulative, recruitment and cumulative vs. recruitment genotype typing resolution scores over time, respectively. Cumulative score for each year is obtained by averaging scores for all samples that are added to the Registry that year or prior to that year, while recruitment score is obtained by averaging scores for only those samples that are added to the Registry in that year.

## 1.4 Discussion

In this paper, we introduced a new measure typing resolution score, TRS, for measuring ambiguity in HLA genotype data. We demonstrated the simplicity and effectiveness of the score by applying it to the data in the Be The Match Registry in order to evaluate the ambiguity in the Registry today and over time.

Typing resolution score has a natural interpretation when it comes to HLA matching. Namely, it describes the probability of a self-match in HLA registry, or the probability of a match between two samples having the same HLA typing. Probability of matching between two typings in the registry is computed as a summation of squared likelihoods of all overlapping imputed genotypes. An HLA typing when matched against itself would result in a completely overlapping list of imputed genotypes and its self-match probability would be equal to its typing resolution score.

Using the proposed self-match based metric, our results illustrate that new recruitment HLA typing performed for the US registry now has very little ambiguity under the current standard of matching for amino acids within the ARS of HLA proteins at HLA-A, C, B, DRB1, and DQB1. In other words, we see that the recruitment typing resolution score approaches a value of 1 for this clinical standard using the current laboratory methods that employ next-generation sequencing or sequence-based typing with panels of group-specific sequencing primers (Figure 3).

For legacy HLA typings such as SSO, we observed significant variation in TRS among populations. LD patterns, HLA diversity of the population group, and the size of the population sample used to generate haplotype frequencies used in TRS computations, all play a significant role in the level of ambiguity. African American samples (AFA) with lower LD have lower scores on average than European American samples (CAU), which have higher LD on average, and higher ambiguity scores (Figure 1a). In addition, the higher HLA diversity of African samples lead to higher ambiguity and lower scores, while the relatively low HLA diversity in European samples lead to higher scores. Asian/Pacific Islander (API) samples are either a mix of multiple population subcategories (Vietnamese, Korean, Japanese, etc.) or donors who joined the registry when the only option was the broad category “Asian or Pacific Islander” which have relatively low HLA diversity within the subgroup, but mixed together constitute a more diverse group. Finally, historical efforts to improve typing methods to separate alleles for the majority group (European Americans) have resulted in a greater reduction of ambiguity in this population (Figure 1(a)).

While the newer sequencing technologies have achieved a great reduction at allelic ambiguity at HLA loci, our analysis shows the need for technologies for experimental resolution of phase, which will lead to a greater reduction of phase ambiguity from what is currently obtainable (Figure 4). HLA phase matching may become more prevalent in the future due to its association with better outcome post-transplant [10], and as such is important to evaluate. While we provide scores for haplotype phase matching on top of high resolution allele matching, haplotype phase cannot currently be experimentally determined without family pedigree information. Therefore, even if clinicians wanted to match haplotypes, they cannot confirm that donor and patient were indeed haplotype matched. The lack of ability to use haplotype phase matching for donor selection, as well as a critical need for confirmation studies of this effect has led to little in the way of clinical uptake.

With high resolution recruitment typing now implemented at these five HLA loci, there is little possibility for improvement in terms of the TRS evaluation method we described. However, we expect that clinical standards for transplant matching will continue to evolve over time, and the TRS metrics can evolve in tandem with these matching standards. Recent publications have suggested a role for additional HLA loci such as DPB1 in transplant outcome [12]. If typing resolution score considered the DPB1 locus, many donors with high scores under the current 5-locus metric would have their overall genotype typing resolution score lowered because of the requirement to impute the alleles at a new untyped locus. On the other hand, per-locus typing resolution scores would remain unchanged. Non-HLA polymorphisms within the major histocompatibility complex (MHC) have also been identified by Petersdorf et al [13]. If haplotype information for both the non-HLA polymorphisms and HLA alleles were known, it would become possible to use the linkage disequilibrium information to extend the TRS metric to include ambiguity for clinically-relevant non-HLA polymorphisms in the MHC. Similarly, polymorphisms outside of the ARS have not been comprehensively evaluated for their impact on transplant outcome. As full HLA gene sequencing proceeds, the number of new HLA alleles in IMGT/HLA [14] will again see a jump as regions of the gene outside of the ARS are systematically interrogated for the first time. In other words, the definition of allele-level matching will also remain a moving target.

KIR polymorphisms have also been identified as determinants of transplant outcome [15,16]. The KIR system has both structural variation in gene copy number and allelic variation [17]. HLA-C alleles and a proportion of HLA-A and HLA-B alleles are ligands for inhibitory KIR genes. As donor selection practices evolve, it may become important to predict the interactions between more probable HLA and KIR polymorphisms. Because KIR and HLA reside on different chromosomes, it is not possible to use linkage disequilibrium information to predict HLA from KIR or vice versa. However, KIR gene frequency information could be used to make predictions when KIR typing is absent. KIR alleles could also be predicted from presence-absence KIR typing data if allele-level KIR haplotype frequencies became available in the future. TRS could therefore be extended to incorporate ambiguity in the KIR region.

While the impact of the HLA typing resolution on unrelated donor selection was the main focus of our analysis, higher-resolution HLA typing data also improves haplotype frequency estimation using the expectation-maximization algorithm [18], patient match likelihood projections when searching the registry [19], and models attempting to identify geographical regions to target donor recruitment in order to remove gaps in HLA coverage [20].

As an attempt to make the method available to the HLA community, we implemented the typing resolution score tool as a part of an existing web-based application HaploStats^1^ developed and maintained by our group. HaploStats is a web application provided by the NMDP Bioinformatics Research group as a tool for the analysis of HLA typing, and to facilitate access to HLA haplotype frequency information relative to various global, country and ethnically specific populations. The snapshot of the HaploStats data input page is shown in Figure 1s. For more information about HaploStats, visit www.haplostats.org.

In conclusion, we present a self-match based typing resolution score metric which has application for comparing HLA typing ambiguity across populations, among registries, varying typing methods, and for individual typings. The results chart the improvement in HLA typing within the Be The Match Registry of the United States from the initiation of DNA-based HLA typing to the current state of high-resolution typing using next-generation sequencing technologies.

## Acknowledgments

This work was supported by Office of Naval Research (ONR) grant N00014-14-1–0848. The content is solely the responsibility of the authors and does not necessarily represent the official views of the Office of Naval Research, the Department of the Navy, the Department of Defense, or the US Government.

## Supplemental Material

**Table 1s.** **Average ambiguity scores for overall and phased genotypes**. This table shows average ambiguity scores for overall genotypes and phased genotypes averaged over all donors and cord blood units for each population separately.

|  | Unphased<br>Genotype | Phased<br>Genotype |
| --- | --- | --- |
| **AFA** | 0.52 | 0.44 |
| **API** | 0.58 | 0.50 |
| **CAU** | 0.73 | 0.59 |
| **HIS** | 0.63 | 0.57 |
| **NAM** | 0.61 | 0.55 |

**Figure 1s.**
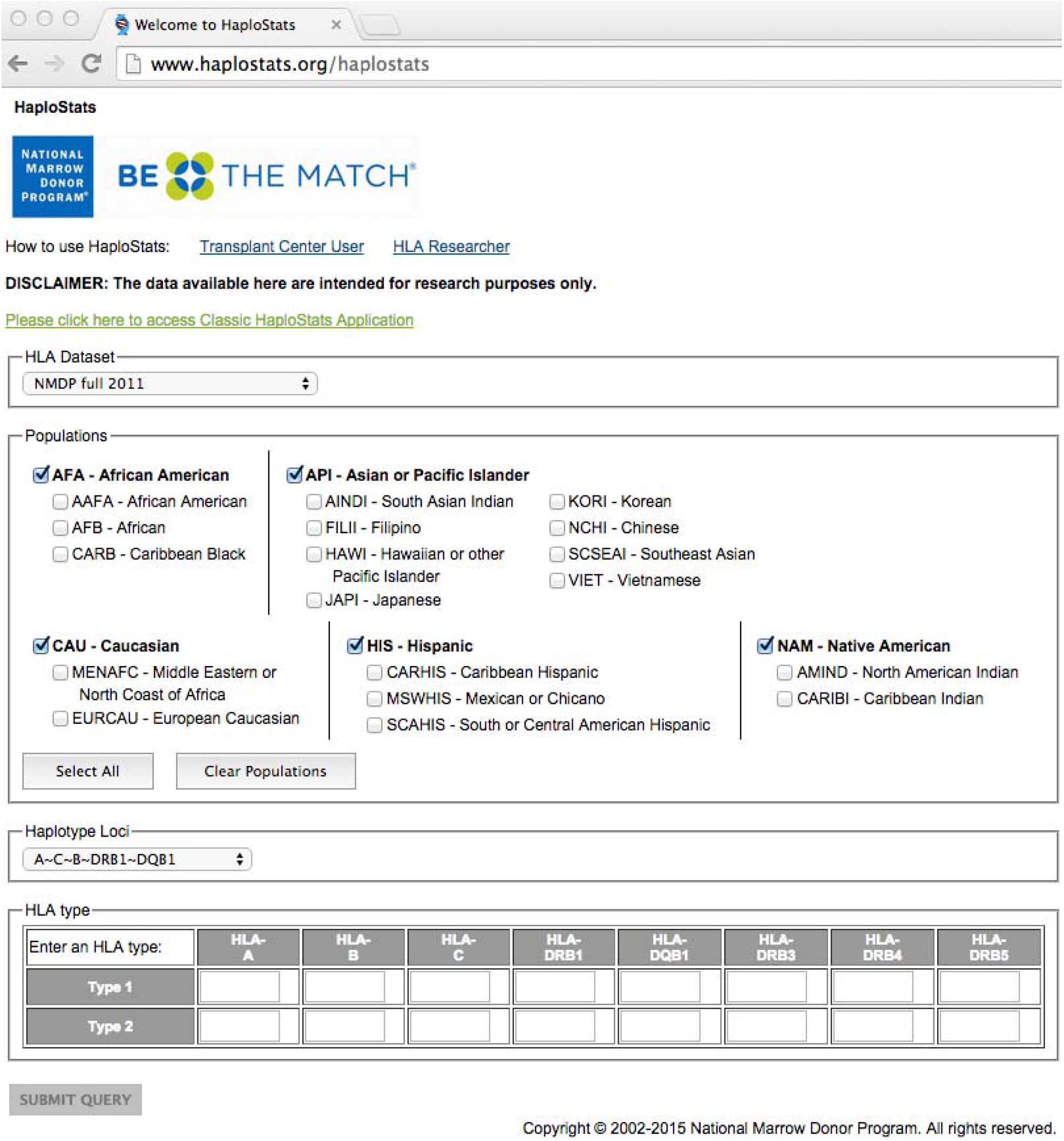
**HaploStats application home page**. This figure shows the snapshot of the HaploStats web-tool home page, where the user can specify the input to the tool which consists of the HLA dataset (HLA haplotype frequencies), populations, the HLA loci that the user wishes to retrieve, and an HLA typing the user wishes to analyze.

**Figure 2s.**
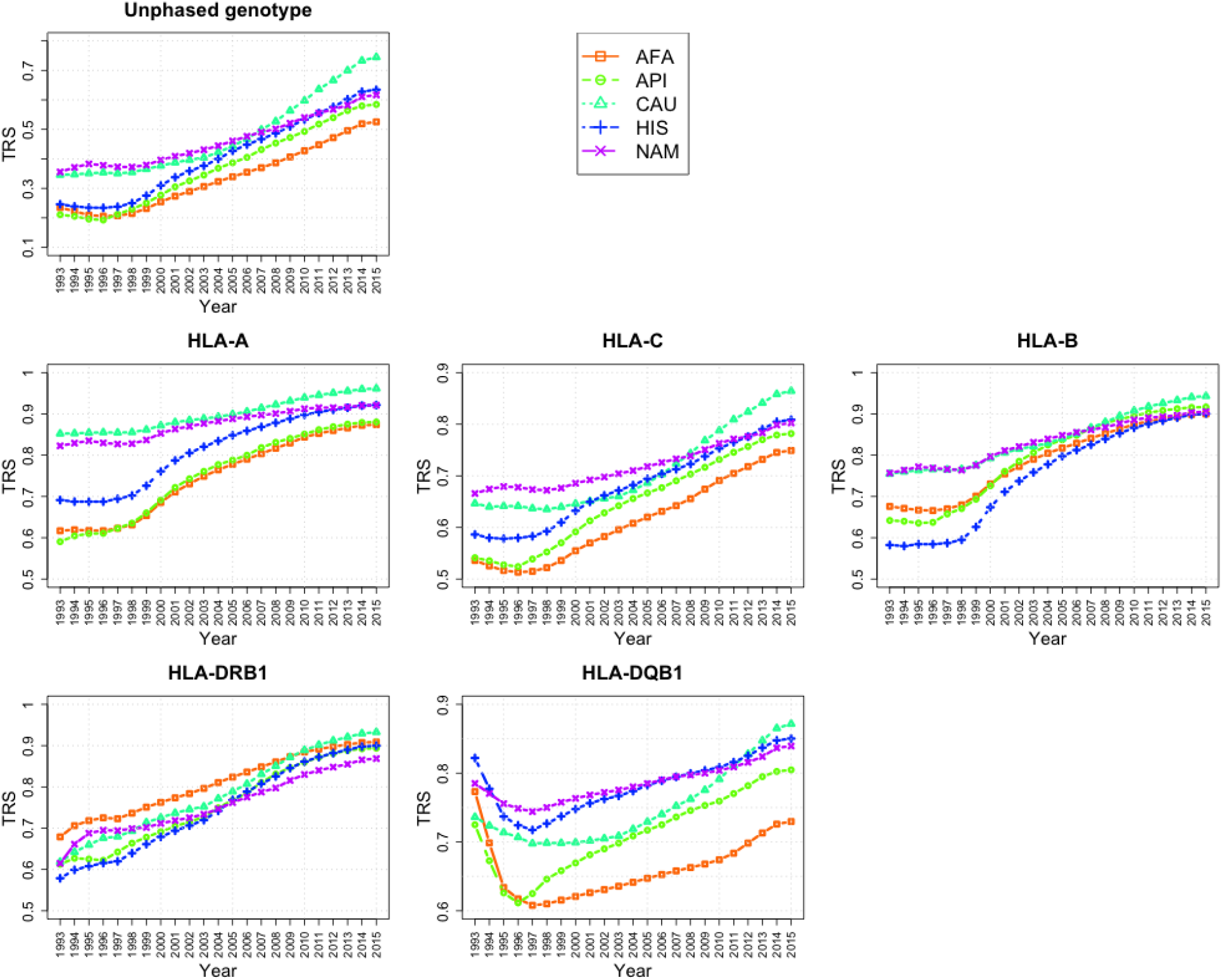
**Cumulative score over time for each population separately**. This figure shows cumulative scores over time for overall genotypes and each HLA locus for each population separately. Cumulative score for each year is obtained by averaging scores for all samples that are added to the Registry that year or prior to that year.

**Figure 3s.**
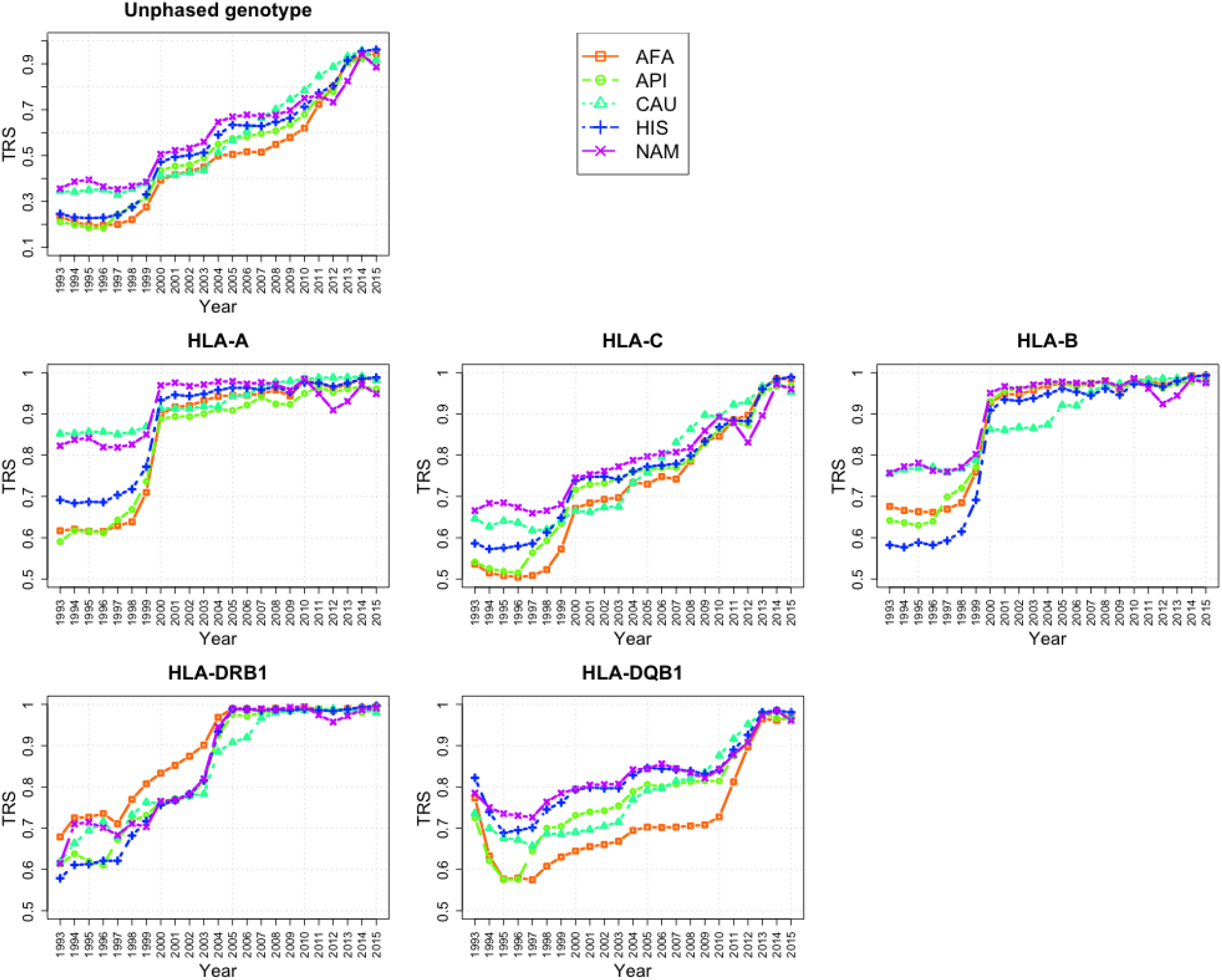
**Recruitment score over time for each population separately**. This figure shows recruitment scores over time for overall genotypes and each HLA locus for each population separately. Recruitment score is obtained by averaging scores for only those samples that are added to the Registry in that year.

## Footnotes

1 http://www.haplostats.org

